# Scalable latent-factor models applied to single-cell RNA-seq data separate biological drivers from confounding effects

**DOI:** 10.1101/087775

**Authors:** Florian Buettner, Naruemon Pratanwanich, John C. Marioni, Oliver Stegle

**Author notes:** Correspondence should be addressed to Florian Buettner, John Marioni or Oliver Stegle.

## Abstract

Single-cell RNA-sequencing (scRNA-seq) allows heterogeneity in gene expression levels to be studied in large populations of cells. Such heterogeneity can arise from both technical and biological factors, thus making decomposing sources of variation extremely difficult. We here describe a computationally efficient model that uses prior pathway annotation to guide inference of the biological drivers underpinning the heterogeneity. Moreover, we jointly update and improve gene set annotation and infer factors explaining variability that fall outside the existing annotation. We validate our method using simulations, which demonstrate both its accuracy and its ability to scale to large datasets with up to 100,000 cells. Moreover, through applications to real data we show that our model can robustly decompose scRNA-seq datasets into interpretable components and facilitate the identification of novel sub-populations.

## Introduction

Single-cell RNA-sequencing (scRNA-seq) is an established tool for assaying variability in gene expression levels between cells drawn from a population. Cell-to-cell differences in gene expression can be driven by both observed and unmeasured factors, including technical effects such as batch, or biological drivers including cell type specific features, such as stage of T cell differentiation ^1^. Importantly, such technical and biological factors can act upon the same genes ^2, 3^, meaning that they should be modeled jointly to fully understand heterogeneity in scRNA-seq data.

Well-established approaches exist for handling observed factors, such as batch or experimental covariates ^4, 5^. Additionally, methods based on singular-value decomposition (SVD) and linear mixed models have been developed in order to capture unwanted variability due to unmeasured factors, first for conventional ensemble RNA-profiling experiments ^6–8^ and more recently for scRNA-seq ^2^. Furthermore, by using informative marker gene sets, methods based on SVD and regression have also been employed to reconstruct cell states, such as the cell cycle or differentiation stages ^2, 9^. More recently, Fan et al. ^10^ introduced PAGODA, a PCA-based method that tests for coordinated overdispersion of sets of genes. However, these existing methods do not model errors in how gene sets are defined, and, more importantly, they fit individual processes independent from each other and do not explicitly account for either the presence of additional unannotated biological factors or confounding sources of variation. Finally, existing factor methods were motivated by relatively small single-cell RNA-seq datasets. Thanks to recent technological advances, it is now possible to generate single-cell RNA-seq datasets containing hundreds of thousands of cells, which requires computationally more efficient methods.

## Results

To address this, we here propose a factorial single-cell latent variable model (f-scLVM) that can capture three sources of variation: i) variation in expression attributable to pre-annotated gene sets; ii) variation attributable to sparse, putatively biologically meaningful, but unannotated gene sets; and iii) variation explained by confounding factors that are expected to affect the expression profile of the majority of genes.

If a factor explains variation in the data, we assume that the expression levels of all genes assigned to it co-vary in a consistent manner, thereby allowing the activity of this factor to be inferred from the data. For annotated factors, we incorporate prior annotations derived from publicly available resources such as MSigDB ^11^ or REACTOME ^12^ into our model, thereby assigning particular biologically related sets of genes to the same factor. The assignment of genes to these sets is refined in a data-driven manner, assuming that only a small number of changes occur (i.e., that the initial annotation is reasonably accurate). For unannotated, but biologically meaningful gene sets, we assume that they contain a small number of genes. Finally, to infer confounding factors we build on the assumption that such factors will have global effects on the expression of large numbers of genes, similar to principles applied in population genomics ^6, 7^.

As well as updating the assignment of gene sets, our model also infers, for a given data set, which factors are most relevant. Inference of model parameters, including gene assignments and factor weights, are determined using computationally efficient variational Bayesian inference, which scales linearly in the number of cells and genes. The posterior distributions over model parameters facilitate a wide range of downstream analyses, including identification of biological drivers, data visualization, the refinement of gene sets and the estimation of residual dataset (Fig. 1a).

**Figure 1.**
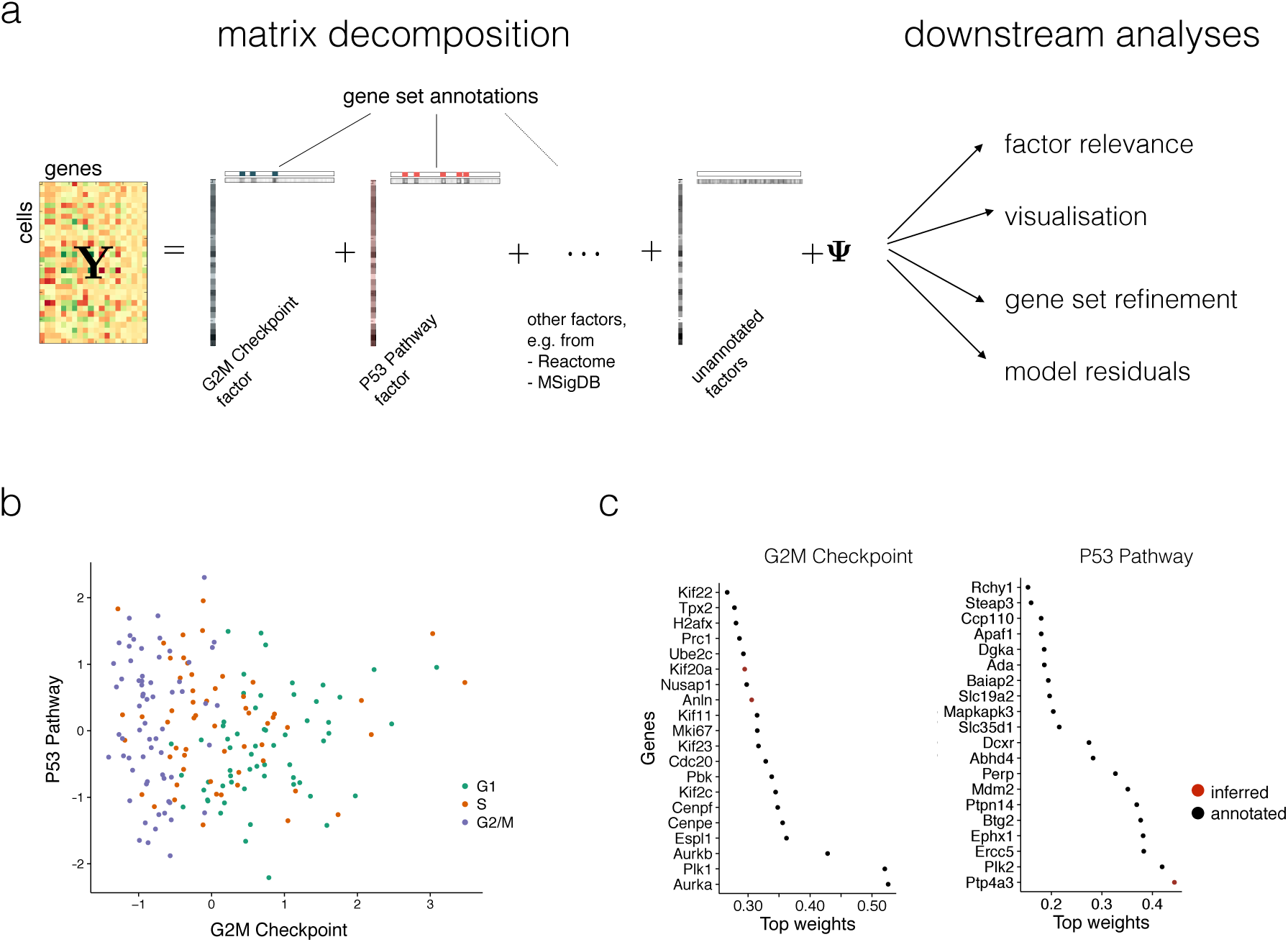
Factorial single-cell latent variable model: approach and motivation. **(a)** f-scLVM decomposes the matrix of single-cell gene expression profiles into factors and weights. Gene sets from pathway databases are used to annotate a subset of factors, with the remainder allowing the existence of unannotated factors. The fitted model can be used for different downstream analyses, including i) identification of biological drivers; ii) visualization of cell states; iii) data-driven adjustment of gene sets and iv) adjustment of confounding factors. **(b)** Bivariate visualization of 182 mouse ES cells, experimentally staged for the cell cycle, using the G2M checkpoint and P53 pathway factors. The inferred G2M checkpoint factor discriminates cells in G2/M phase from the remaining cell population. **(c)** Weights for the most important genes in the P53 pathways and G2M checkpoint factors, showing both genes that were pre-annotated by MSIGDB (black), and genes added by the model (red).

As a first illustration, we applied f-scLVM to a dataset of 182 mouse embryonic stem cell (mESC) transcriptomes, where each cell was experimentally staged according to its position within the cell cycle ^2^. Consequently, across the entire population, we expect that the cell cycle is the major source of variation. Indeed, when applying f-scLVM using 44 core molecular pathways derived from MSigDB ^11^, the method robustly identified four factors, including G2/M checkpoint and P53 pathway (Supp. Fig. 1).

These two factors could be used to stratify the cells by their position in the cell cycle (Fig. 1b). Other methods, including PAGODA, inferred a larger number of collinear and partially redundant factors that less accurately discriminated cells by cell cycle stage (Supp. Fig. 2), underscoring the importance of jointly modeling all annotated and unannotated factors. Furthermore, unlike existing methodology, f-scLVM allowed for data driven refinement of gene set annotations (Fig. 1c). We observed that the model modified the G2/M checkpoint factor by adding two genes, Anln and Kif20a, both of which are well-characterized cell cycle regulators ^13, 14^. Similarly, the model identified Ptp4a3, a known target of P53 ^15^, as an additional member of the P53 pathway factor.

Given these promising results, we next used simulated data to assess how robustly our model can identify relevant factors and complete gene set annotations. Over a wide range of simulations, where we varied the number of annotated factors and simulated confounders, the degree of overlap between gene sets, the number of cells and the size of the annotated gene sets, we consistently observed that f-scLVM more accurately identified the true simulated drivers than other factor analysis methods (Fig. 2a and Supp. Fig. 3a-e). Our method showed the greatest improvement in performance over existing methods when multiple unannotated factors were simulated and when the gene sets explaining the most variance in the data contained a substantial number of overlapping genes (Fig. 2b). We also confirmed that f-scLVM was robust to extremely sparse datasets, typical of droplet-based approaches (Supp. Fig. 3f,g).

**Figure 2.**
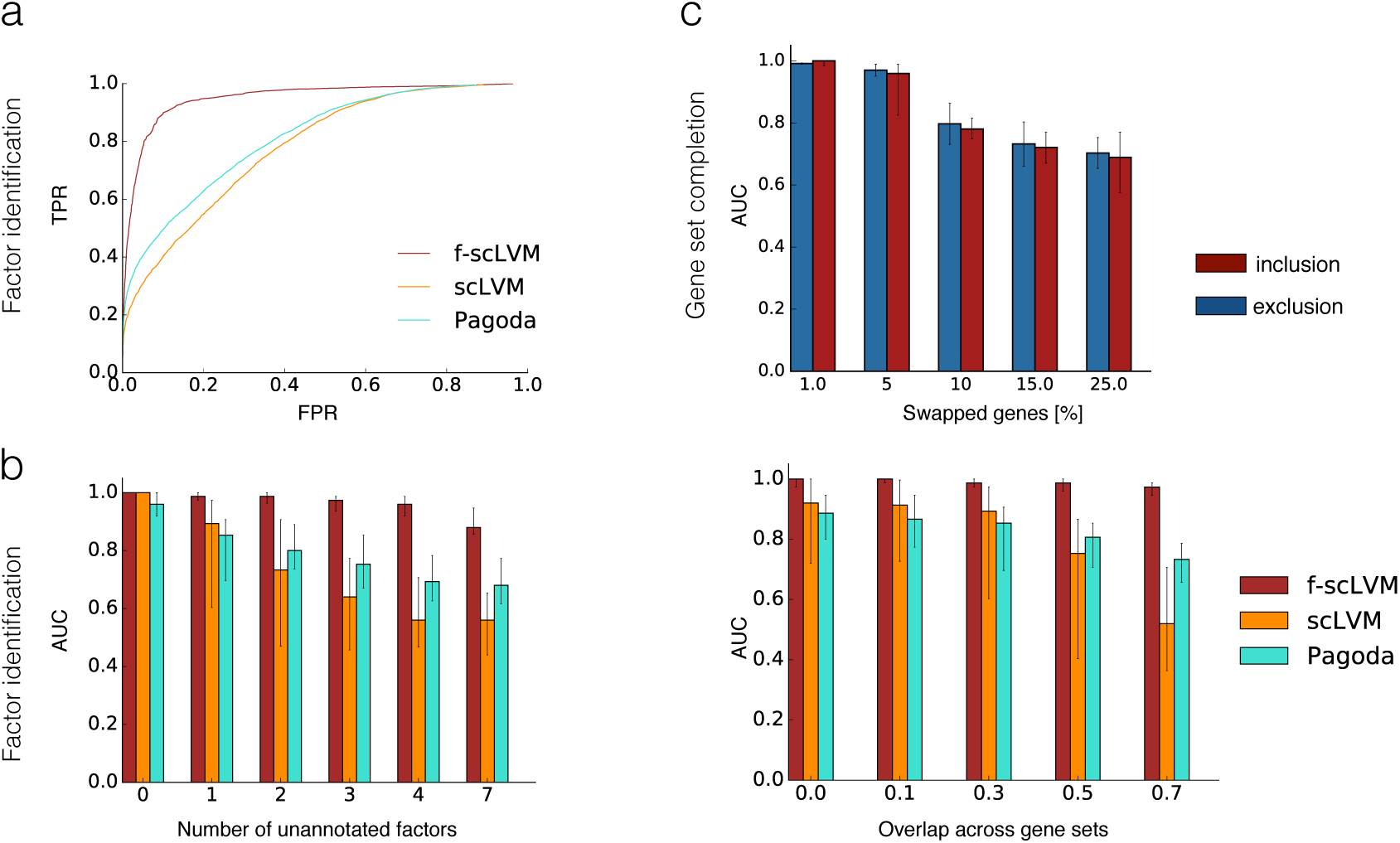
Model validation using simulated data. **(a,b)** Accuracy of f-scLVM and alternative methods for recovering the set of simulated drivers of gene expression heterogeneity. **(a)** Receiver operating characteristics (ROC) pooled across different simulated datasets (see Supp. Table 1). **(b)** Area under the ROC curve (AUC) when simulating an increasing number of unannotated factors not included in the pathway database (left), and when considering increasing overlap between simulated gene sets (right). **(c)** Ability of f-scLVM to augment gene sets when an increasing proportion of genes in the annotation were falsely assigned. Shown is an AUC for correctly including genes omitted from gene sets (red) and for removing genes that were incorrectly annotated (blue). Bar plots in **(b,c)** show the median AUC across 50 repeat experiments per settings with error bars corresponding to 25% and 75% percentiles.

Finally, we considered datasets with simulated errors in the gene set annotation to assess the model’s ability to adjust gene sets, a feature that is unique to f-scLVM (Methods). We observed that the model accurately identified genes that should be excluded from and added to gene sets (Fig. 2c and Supp. Fig. 4). Unsurprisingly, as the fraction of errors in the gene set annotation increased, the ability of the model to recover the true sets declined – however, in the more realistic setting where only a small fraction of genes were poorly annotated (1-10%), our model performed extremely well.

Having evaluated our approach, we next applied f-scLVM to a population of 3,005 neuronal cells ^16^. Using gene sets derived from REACTOME pathways as annotation, our model supported the importance of a set of factors similar to those identified by methods such as PAGODA (Fig. 3a), but with important differences (e.g. Innate Immune System; Supp. Fig. 5). Additionally, our model suggested a refined annotation for some of the most important gene sets identified (Fig. 3a), with, on average, 10% of genes being added and 3% of genes being removed for the top 20 annotated factors. Furthermore, the model identified unannotated factors with a high relevance score (Fig. 3b), demonstrating the importance of modelling such factors. We observed that many of these unannotated factors were sparse and captured differences between cell types that are not readily reflected in the pathway annotations (Supp. Fig. 5,6a-c).

**Figure 3.**
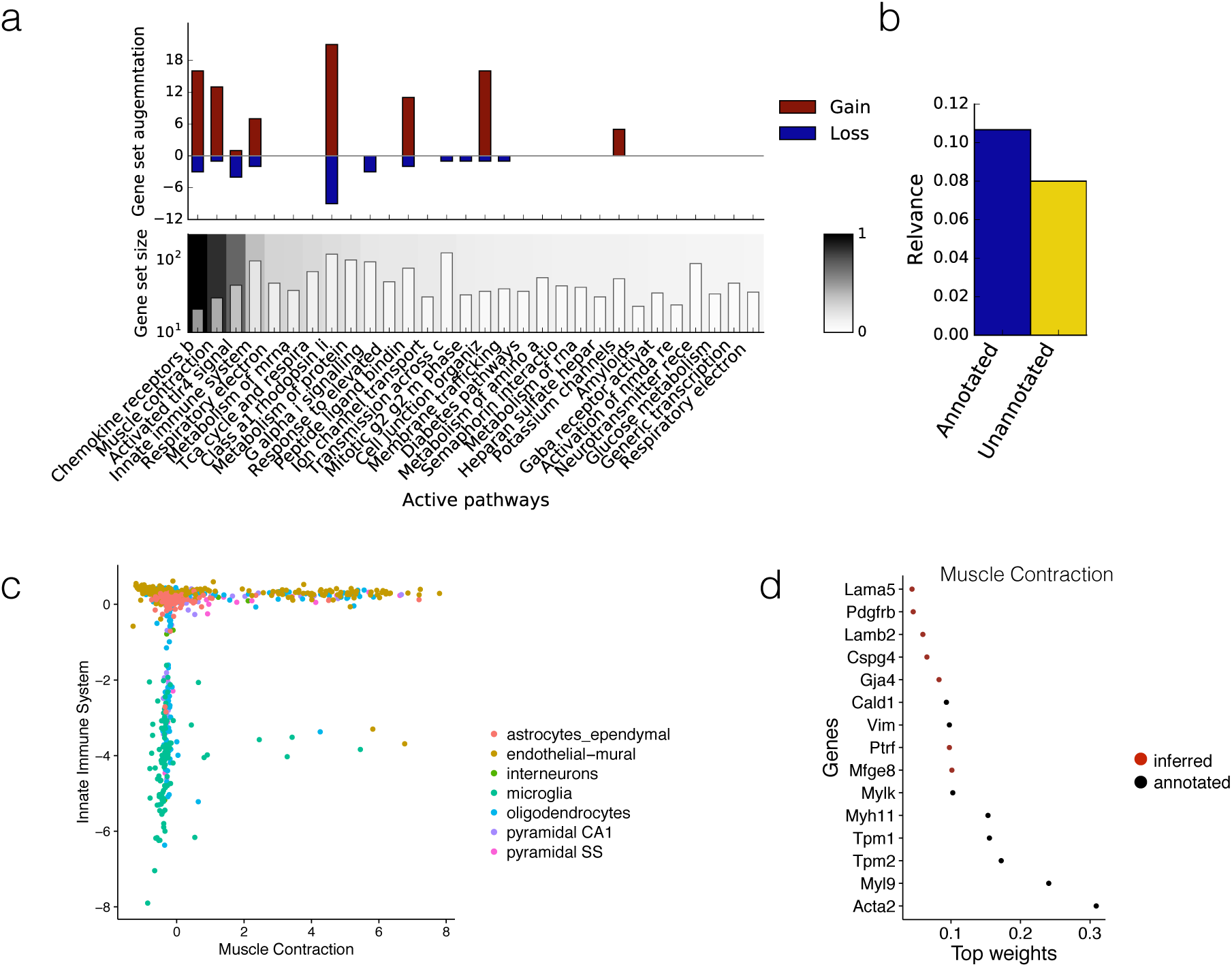
Application of f-scLVM to neuronal cells. **(a)** Factor relevance and gene set augmentation for the most important 30 factors identified by f-scLVM based on REACTOME pathways. Bottom panel: Identified factors and corresponding gene set size ordered by relevance (white = low relevance; black = high relevance). Top panel: Gene set augmentation, showing the number of genes added (red) and removed (blue) by the model for each factor. **(b)** Breakdown of the cumulative factor relevance of annotated and unannotated factors (see also Supp. Fig. 5). **(c)** Bivariate visualization of cells using the factors muscle contraction and innate immune system. Colors correspond to cell types identified in ^16^. **(d)** Weights for the most important genes in the muscle contraction factor, showing both genes that were pre-annotated by REACTOME (black), and genes added by the model (red).

The top ranked annotated processes separated the cells into well-defined groups, with the endothelial-mural cells being stratified into two populations by the muscle contraction factor (Fig. 3c). Notably, our model augmented the corresponding gene set by activating several genes that have previously been implicated in muscle contraction but that were not present in the pre-defined REACTOME gene set (Fig. 3d). Among the 13 identified genes were several known markers of vascular smooth muscle cells, including Rgs4, Mtfge8, and Notch3 ^17–20^. A second major driver identified by f-scLVM was the innate immune system factor, which, in particular, separated microglia (nervous system immune cells) from the remaining cell types (Fig. 3c). Moreover, similar to the muscle contraction factor, the gene set was also augmented with meaningful genes (Supp. Fig. 6d).

In addition to the populations of neurons characterized by Zeisel et al., we also applied f-scLVM to a variety of other existing datasets, identifying relevant factors that explained complementary axes of variation, augmenting gene sets and observing that a considerable proportion of the variance explained could be attributed to unannotated factors (Supp. Analyses) and we assessed the robustness of the annotated factors identified by f-scLVM (Supp. Fig. 10).

Finally, given the increasing trend to generate scRNA-seq datasets containing tens to hundreds of thousands of cells, we contrasted the computational efficiency of our model with a variety of factor analysis models (Fig. 4a, Supp. Fig. 7). We observed that irrespective of the number of cells, f-scLVM had a lower runtime than other approaches, with a linear scaling time in the number of cells as opposed to quadratic or even cubic relationships for other approaches. As a consequence, f-scLVM can be applied to interrogate large-scale droplet-derived datasets.

**Figure 4.**
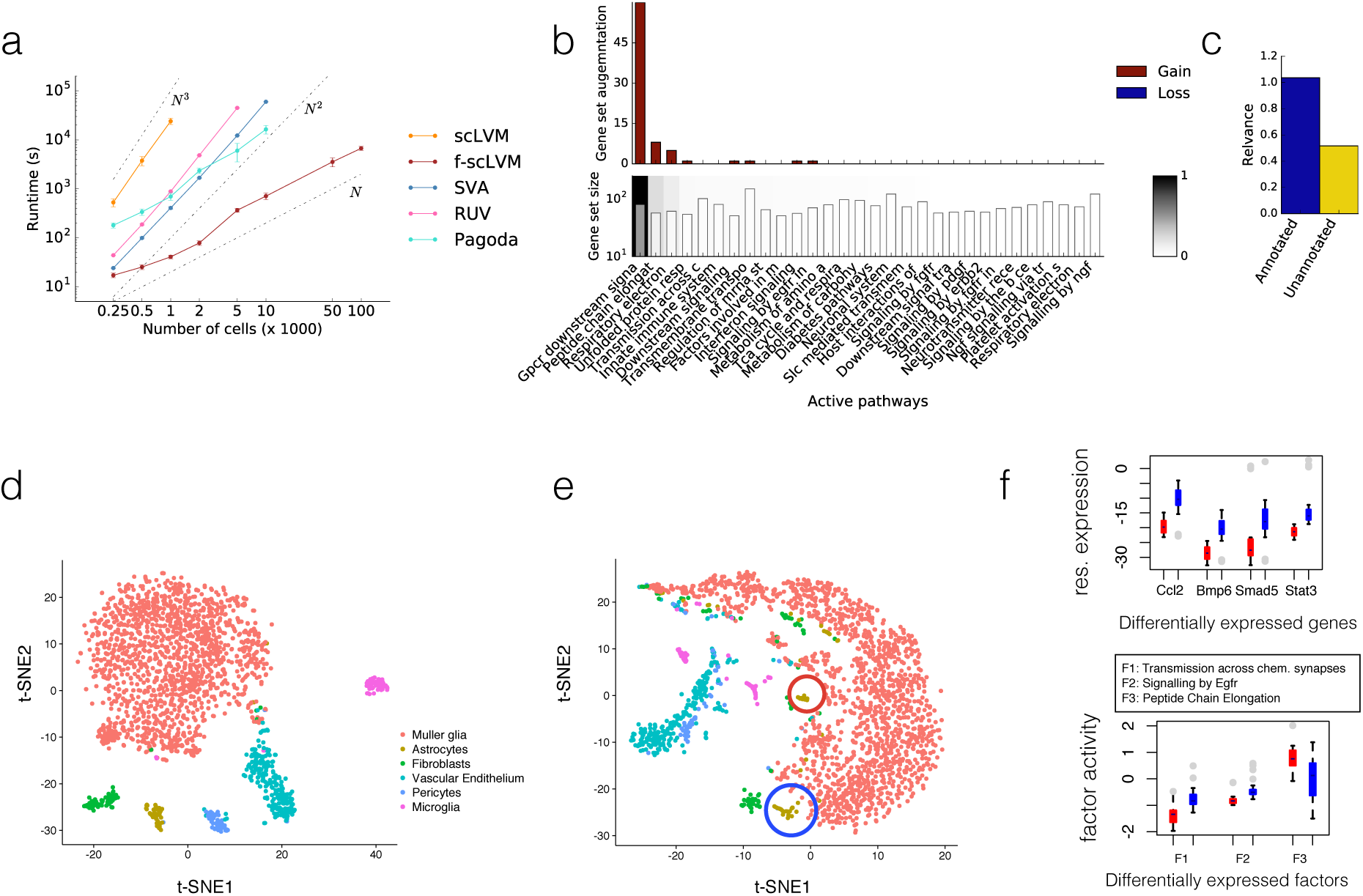
Application of f-scLVM to large-scale scRNA-seq datasets. **(a)** Scalability of f-scLVM, showing is the empirical runtime when applying f-scLVM and alternative factor models (RUV, SVA, scLVM, PAGODA) to datasets with increasing size. f-scLVM scales linearly in the number of cells, allowing its use on large dataset with up to 100,000 cells. None of the existing methods could be applied to the largest dataset. **(b-f)** Application of f-scLVM to 49,300 retina cells profiled using Drop-Seq. **(b)** Factor relevance and gene set augmentation for the most important 30 factors identified by f-scLVM based on REACTOME pathways. Bottom panel: Identified factors and corresponding gene set size ordered by relevance (white = low relevance; black = high relevance). Top panel: Gene set augmentation, showing the number of genes added (red) and removed (blue) by the model for each factor. **(c)** Breakdown of the cumulative gene expression variance explained by annotated factors and unannotated factors. **(d)** Visualization of a subset of 2,145 cells using a non-linear t-SNE embedding. Colors correspond to cell types identified in ^21^. **(e)** Analogous t-SNE embedding as in **(d)**, however on residual data (Methods, see also Supp. Fig 8). The analysis on the residual dataset revealed additional sub structure between cells, including a new sub population of astrocytes. **(f)** Genes and factors that were differentially expressed (FDR < 10%) between the newly-identified astrocyte clusters highlighted in **(e)**. Grey dots denote outlying cells.

To illustrate this, we applied f-scLVM to expression profiles from 49,300 retinal cells profiled using Drop-seq ^21^. Again, our model identified biologically plausible processes, including GPCR signaling and transmission across chemical synapses, as explaining variability in the data (Fig. 4b). As previously, unannotated factors explained substantial variation in the data (Fig. 4c). In this case, this variation could be attributed to a single factor, which affected a large number of genes (Supp. Fig. 8a), suggesting that it may explain confounding effects. Indeed, this factor was correlated with the cellular detection rate (Supp. Fig 8f), a known confounding feature of scRNAseq datasets ^3^.

Given this, we considered the residual dataset generated after regressing out the effect of this factor (Methods, Supp. Table 2). When focusing on a set of six related and well defined cell types identified in the primary analysis – Müller glia, astrocytes, fibroblasts, vascular epithelium, pericytes and microglia – the residual data revealed two sub populations of astrocytes (Fig. 4d,e), as well as two subpopulations of microglia (Supp. Fig. 8b-d) that are not observed in the unadjusted data. In total, 1,024 genes were differentially expressed between the identified astrocyte populations (FDR<10%; Supp. Table 3) and these were enriched for processes related to immune response and activation of astrocytes, such as inflammatory response, BMP signaling pathway, and cellular response to BMP (Supp. Table 4), and included known genes related to reactive/inflammatory processes in astrocytes such as Ccl2 ^22^. In addition BMP signalling is known to activate distinct downstream transcription factors, including Stat3 and Smad5, which show the expected pattern of behavior between the two newly-identified populations ^23^ (Fig. 4f). Taken together, these results show that f-scLVM can be used to infer biological and confounding factors from large datasets.

More broadly, we considered a range of available datasets and investigated the nature of unannotated factors inferred by our model. We observed, as above, that these factors were often associated with technical experimental features that have previously been suggested to underpin variability in scRNA-seq data, including the number of expressed genes and sequencing depth (Supp. Fig. 9). However, these associations were often weak, suggesting that the inferred hidden factors help to capture additional unwanted variation that cannot be assigned to measured covariates.

## Discussion

Herein, we have proposed a scalable factor analysis approach to comprehensively model the sources of single-cell transcriptome variability. Unique to our model is the ability to jointly infer both annotated and unannotated factors and to augment predefined gene sets in a data driven manner. Additionally, f-scLVM is computationally efficient, allowing analysis of very large datasets containing hundreds of thousands of cells.

We have validated our model using simulations as well as using real data where the sources of transcriptome variability are well understood. Subsequently we have applied the model to a range of different studies, demonstrating its ability to infer drivers of transcriptome variation, adjust gene sets to discover new marker genes and account for hidden confounding factors in the data.

Of course, our model is not free of limitations. A general challenge for any method is to reliably differentiate confounding factors from biological signal. f-scLVM addresses this through specific assumptions on the effect of these factors (sparse versus dense) in conjunction with leveraging gene set annotation from pathway databases. However, there is no silver bullet solution and it is hence necessary to interpret the model results and, in particular, the unannotated factors in the context of a given dataset.

A second potential caveat is the lack of accurate gene set annotation, which will necessarily impact the quality of the results. To mitigate this challenge, f-scLVM models possible errors in the annotation explicitly and augments gene sets in a data driven manner. However, such inferences have limitations. One important challenge is collinearities between annotated factors and true biological differences. For example, if cells in different cell cycle stages are systematically associated with different cell types, the results from gene set refinements may be misleading and collapse two distinct biological processes into a single factor.

There are other technical aspects of the model that could be improved in the future. The noise model we use at present is based on a Hurdle model ^3^, which could be adapted to more specifically model the noise properties of different experimental platforms. Also, our model is intrinsically linear and assumes that the inferred factors have linear additive effects. An important area of future work will be to explicitly model interactions between factors.

## Methods

Methods and any associated references are available in the online version of the paper and further information is provided in the Supp. Methods.

## Acknowledgements

We thank Martin Hemberg for comments on the manuscript and Bertie Göttgens for helpful discussions. F.B. received support from the UK Medical Research Council (MRC Career Development Award in Biostatistics).

## Author contributions

F.B. and O.S. conceived the method, designed the experiments and analyzed the data. F.B. performed all experiments with help from N.P. F.B. J.C.M. and O.S. interpreted the results and wrote the paper.

## Online Methods

### Code availability

An open source implementation of f-scLVM for reviewing purposes is available at: https://github.com/PMBio/f-scLVM.

### The factorial single-cell latent variable model (f-scLVM)

f-scLVM is based on a variant of matrix factorization, decomposing the observed gene matrix into a sum of sum of contributions from C measured covariates, A annotated factors, whose inference is guided by pathway gene sets, and H additional unannotated factors:

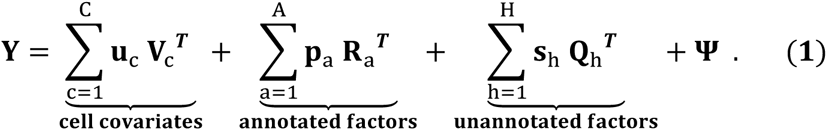

Here, **Y** denotes the gene expression matrix where rows correspond to each of N cells and columns correspond to G genes. The vectors **u**_c_, **p**_a_ and **s**_h_ are known cell covariates, as well as cell states for annotated and unannotated factors, and **V**_c_, **R**_a_ and **Q**_h_ are the corresponding regulatory weights of a given factor on all genes. The matrix **Ψ** denotes residual noise, with its specific form depending on the noise model employed (see below). For the statistical derivation (see also Supp. Methods), we express the model in Eq. (1) using matrix notation, collapsing the factors into a factor activation matrix **X** = [**u**_1_,..,**u**_C_, **r**_1_,.., **r**_A_, **s**_1_, …, **s**_H_] (with the comma denoting concatenation of columns), where each factor is enumerated using an indicator k = 1.. K, and K denotes the total number of fitted factors K = C + A + H. The analogous matrix representation is used for weights **W**, resulting in

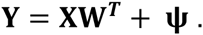

Known covariates, annotated factors and unannotated factors then correspond to different distributional assumptions on the column vectors of the matrices **X** and **W** (Supp. Methods). For brevity, we omit the cell covariates in the derivation, for which the factor states are observed (see Supp. Methods).

Standard multivariate normal prior distributions are used for the factor states of both annotated and unannotated factors. For annotated factors, we employ two levels of regularization on the corresponding parts of the weight matrix **W**. First, gene sets are used to guide a sparsity prior on the rows of **W** ^24^, thereby confining the inferred weights to the set of genes annotated in the pathway database. A second level of regularization is then used to achieve sparseness on the level of factors, allowing the model to deactivate factors that are not needed to explain variation in the data.

#### A regulatory sparseness prior for linking factors to biological processes

Gene set annotations are used to inform a spike and slab prior on the elements of **W**. The regulatory weight of factor *k* on gene *g* is modeled using a mixture of a Normal distribution (with factor-specific precision *α*_*k*_), if the regulatory link is active, and a delta distribution to force the weights of inactive regulatory link to zero

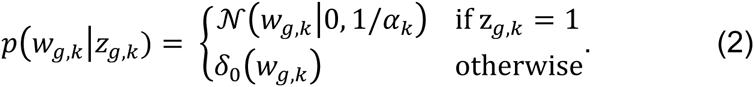

The indicator variable *z*_*g,k*_, which determines whether factor k regulates gene g, is unobserved. However, annotations derived from pathway databases provide additional evidence for inferring *z*_*g,k*_ and for obtaining interpretable factors

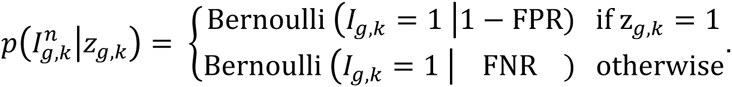

Here, 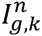 is a binary variable that determines whether gene *g* is annotated to pathway *k* in the annotation database. The pathway annotation is observed for each cell *n*, thereby scaling the evidence from the annotation and the expression data irrespective of dataset sizes (Supp. Methods). The rate parameter FPR corresponds to the probability of a false positive (incorrect) assignment and FNR denotes the probability of a false negative (missing) assignments. In the experiments, we use FNR = 0.001 and FPR = 0.01 and employ an uninformative prior on the indicator variables, *z*_*g,k*_ ~Bernoulli (0.5).

During model training, the posterior distribution of the indicator variables *z*_*g,k*_ is then inferred from the observed expression data and the prior annotation jointly. This approach allows gene set annotations to be incorporated as well as data-driven augmentation. Once trained, the marginal posterior estimates of *z*_*g,k*_ can be used to identify genes that are added to or removed from a particular process.

#### Identifying expression drivers using automatic relevance determination

In addition to sparseness of regulatory effects, f-scLVM employs a second level of regularization using automatic relevance determination (ARD) ^25^. The ARD prior is widely used in (probabilistic) matrix factorization models to deactivate factors that are not needed. This is achieved by placing a hierarchical prior on the precision of the normal priors for active links (Eqn. (2)) *α*_*k*_ ~ Γ(*a*, *b*). The parameter *α*_*k*_ will be large for factors with low relevance, which corresponds to low prior variance, thereby driving the regulatory weights to zero. The prior variance 1/*α*_*k*_ can also be interpreted as a measure of the regulatory impact of a factor and corresponds to the expected variance explained by the factor, for the subset of genes with a regulatory effect (see downstream analyses).

#### Modeling unannotated factors

In addition to annotated factors, f-scLVM estimates the effect of a fixed number of unannotated factors jointly. In the experiments, we consider two types of unnoted factors. First, to infer likely confounding factors, we build on the insight that confounders tend to have broad effects and regulate larger sets of genes, a principle that is widely used in population genomics ^6, 7^. This prior belief is encoded using the Bernoulli prior *z*_*g,k*_ ~Bernoulli(0.99), and hence the weights for these factors are effectively only regularized by the ARD prior. Optionally, f-scLVM can also be used to infer an additional set of sparse unannotated factors. These factors can, for example, be used to model additional biological variation that is not well captured by the factors in the annotation. These sparse actors are modeled using a Bernoulli prior that favors a small number of active links *z*_*g,k*_ ~Bernoulli(0.01). The decision, which model to train, can be guided by heuristics and diagnostics; see section *Diagonistics and f-scLVM parameter settings* for details on the selection of specific models.

#### Noise model

f-scLVM supports alternative noise models to accommodate different RNA-sequencing protocols. First, a standard option is the lognormal noise model, where the expression matrix ***Y*** consists of log count values which are modeled assuming iid heteroscedastic residuals **Ψ** (Supp. Methods). Modeling different residual variances for each dimension (gene) helps to account for varying extent of over dispersion and allows the model deactivating some input dimensions, an approach widely adopted in conventional factor analysis ^26^.

In order to model the zero inflation resulting from prominent dropout effects for protocols such as Drop-seq ^21^, f-scLVM can alternatively be run in conjunction with a zero inflation (Hurdle) noise model. A separate Bernoulli observation noise model is used, when no expression (zero count values) is observed for any specific expression value, while all remaining values are modeled using the aforementioned log Gaussian noise model. Formally, we define the factor analysis model on latent variables, **F** = **XW**^T^, and use the compound likelihood:

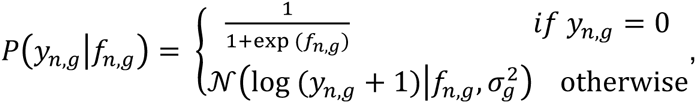

where analogous to the log normal noise model, *y*_*n,g*_ correspond to log count observations. Note that in the absence of zero counts, this noise model is reduced to the basic noise model.

Finally, if zero-inflation is less likely, for example in deeply sequenced datasets with larger quantities of starting material per cell, f-scLVM can also be used in conjunction with a classical Poisson noise model. The inference approach is analogous to the dropout model, however assuming the following likelihood model:

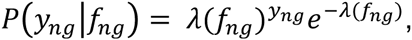

with link function 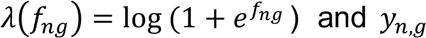 now denoting raw count values.

#### Parameter inference

Closed-form inference in sparse factor analysis is not tractable. In order to achieve scalability to large numbers of cells and genes, we employ deterministic approximate Bayesian inference based on variational methods ^27^. The core idea of variational Bayes is to approximate the true posterior distribution over all unobserved variables using a factorized form. This assumption of (partial) factorization of the posterior allows derivation of an iterative inference scheme, updating posterior distributions for individual parameters in turn, given the state of all others. For full details and the update equations for the f-scLVM see Supp. Methods.

#### Downstream analysis

The fitted f-scLVM model allows for a range of different downstream analyses. Factor relevance: The relevance of annotated pathways factors can be deduced from the ARD variance 1/*α*_*k*_, which corresponds to the expected explained variance of factor *k* for the subset of genes with a regulatory effect (e.g. Fig. 3a,b).

Visualization: The posterior distribution over the inferred factors **X** allows for the visualization of cell states (e.g. Fig. 3c). This is possible both for annotated factors and for unannotated factors, where in particular sparse unannotated factors frequently tend to capture additional structure between cell types (Supp. Fig. 5,6).

Gene set refinement: By comparing the posterior distribution on the indicator variables *z*_*g,k*_ with the prior gene set annotations *I*_*g,k*_, it is possible to identify individual genes that were added to or removed from a pathway factor during inference (e.g. Fig. 3a,d). We use the posterior threshold of 0.5 to identify genet set augmentations in the annotation.

Estimation of residual expression datasets and imputation: The learnt factor **X** in combination with the learnt regulatory weights **W** can also be used to calculate residual dataset with the effect of selected factors removed or to obtain imputed datasets. When using the dropout noise model, expression residuals are estimates based on the latent expression values **F** (see *Noise model* above). In this instance the model implicitly uses the dropout noise model to implicitly impute zero values prior to estimating expression residuals (see Supp. Methods).

#### Relationship to other factor analysis models

Several existing factor analysis methods are related to f-scLVM. First, factor analysis with dense unannotated factors is used to adjust for unwanted variation in bulk datasets, including SVA ^6^, RUV ^8^ and PEER ^7^. However, unlike f-scLVM, these methods do not model annotated factors using gene sets and hence are not designed for identifying biological drivers. Second, methods such as PAGODA ^10^ use gene set annotations to infer interpretable factors. However, this model ignores variation outside the annotation and it infers factors sequentially, which leads to collinearities between factors (Supp. Fig. 2b,e,f). Finally, there exist methods based on sparse factor analysis, including non-parametric methods and factor models that account for the specifics of single-cell transcriptome noise. Again, these methods do not utilize gene set annotations. f-scLVM generalizes many of these methods and in particular offers favorable computational efficiency. For further details and a tabular comparison of the features offered by different methods see Supp. Methods Sec 2.

### Implementation of comparison partners

We compared the performance of f-scLVM to a number of alternative factor models. First, we ran PAGODA using the scde R package ^10^. Briefly, PAGODA infers a gene-specific residual variance by deriving cell-specific error models accounting for dropout effects. This error model is then used to perform weighted PCA ^28^ on individual gene sets in turn, followed by a ranking of gene sets based on the explained variance of the leading PC. We also considered the single-cell latent variable model (scLVM ^2^), which analogously to PAGODA infers independent low-rank factors based on predefined gene sets. The proportion of average variance explained by individual factors for the set of annotated genes, as determined using the variance decomposition described in ^2^, was used to rank factors. A second class of methods we compared to are factor models that do not explicitly incorporate gene set annotations. Among these we used a conventional PCA fitted to the set of all expressed genes. Second, we applied the recently proposed Zero-Inflated Factor Analysis model (ZIFA) ^29^, a factor analysis implementation that explicitly models dropout events. Third, we applied a sparse factor analysis model based on the Indian buffet process (IBP), a non-parametric model that automatically infers the most appropriate number of sparse factors. None of these methods annotates the inferred factors and hence we implemented a post-processing step based on a competitive gene set enrichment to annotate the learnt factors using the same gene sets used to fit f-scLVM. We then used the enrichment p-value to rank the annotated individual gene sets as potential biological drivers.

For runtime assessments, we additionally considered two approaches to account for unwanted variation. SVA timings were reported using the R implementation of the SVA package ^30^, considering a fixed number of surrogate variables that correspond to the true number of simulated factors. RUV runtime results were obtained using the RUV2 function from the R implementation, which estimates and adjusts for unwanted variation using control genes. Runtimes estimates were obtained using the time module in python (time() function) and the proc.time() function in R; all simulations were run on 8 cores of an Intel Xeon 2.60GHz CPU.

### Datasets and preprocessing

#### Diagnostics and f-scLVM parameter settings

By default f-scLVM is fitted using annotated factors guided by gene set annotations and additional dense unannotated factors that capture unwanted variation. However, for some datasets this set of factors may not be sufficient to explain the observed heterogeneity, e.g. because potential differences between cell types may not be well reflected by the provided annotations. In this case, it is advised to infer an additional set of sparse unannotated factors. A suitable diagnostic for this decision are excessive augmentations of the annotated gene sets such that the inferred factor is unlinked to the annotated biological process. In the software implementation of f-scLVM sparse unannotated factors are activated if the standard model changes (gains or losses) at least 100% of annotations for at least one annotated factor. For sparse and dense unannotated factors, we considered five and three factors by default, respectively. Note that because of the ARD prior, the model is robust w.r.t. to the number of dense unannotated factors, provided a sufficiently large number is inferred (Supp. Fig. 3h).

#### Simulation study

We simulated gene expression matrices based on a linear additive model, an assumption that is motivated by the generative model that underlies both f-scLVM and the all existing alternative approaches, all of which are based on variants of linear factor analysis models (Supp. Methods). We simulated effects from between three and ten active pathway factors with partially overlapping gene sets for each factor (see below), additional effects due to unknown confounding factors, and observation noise. We considered a total of 44 simulation settings, considering variable dataset sizes (cell count), variable numbers of active pathway factors and increasing numbers of simulated unannotated confounding factors. Additionally, we varied the overlap of genes annotated to individual pathways, the size of the individual gene sets, and simulated a certain degree of noise in the annotation provided to each respective model, by simulating a certain proportion of false negative/false positive annotations and by swapping genes between active factors (see Supp. Table 1 for full details of simulation parameters).

Each simulated dataset consisted of between 20 - 500 synthetic cells and 6,000 genes. Gene set sizes were determined by sampling from REACTOME pathways (considering 421 pathways with 20 to 933 genes). When ranking active pathways, we compiled an annotation consisting of the true drivers and an additional set of 15 non-active pathways as a negative set, and provided it to each considered method. Confounding factors, if simulated, were generated analogously to the approach described in ^31^, assuming broad effects affecting between 400 and 3,000 randomly selected genes. The annotations of simulated pathways were generated sequentially, ordering pathways by decreasing size and drawing genes with a selected overlap to already existing pathways. Pathways were simulated to have overlapping gene sets, between 0.0 (no overlap) and 0.7 (70% of the gens overlap). To test for the impact of the size of gene sets, we additionally considered sampling REACTOME pathways with between 20-50, 50-100, and 100-200 genes. To assess the robustness of f-scLVM to incorrect gene set annotations, we simulated between 1% and 50% of false negative and between 1% and 10% false positive genes in the gene set annotations of individual factors. We also considered more challenging miss-annotations by introducing gene-swaps between pairs of active factors (for between 1% and 25% of all genes). Factor activations as well as non-zero regulatory weights were drawn assuming a unit variance normal distribution. Residual noise was simulated as normally distributed with standard deviation 0.1. When dropout was simulated, we considered two alternative dropout mechanisms. First, we model a threshold effect by setting all values less than a given threshold to 0; this reflects a limit of detection where small numbers of molecules cannot be detected reliably. Second, we considered modeling the probability of dropout events as a function of the true expression level, assuming an exponential relationship ^29^: 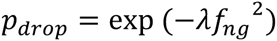, with *f*_*n,g*_ being the latent expression level introduced above and *λ* the exponential decay parameter. Both dropout processes are simulated, where each setting is parameterized by *λ* and the threshold value, which corresponds to the lower limit of detection. For each simulation setting, 50 independent datasets were generated.

To assess the performance of f-scLVM and alternative methods, we consider the receiver operator characteristics (ROC) for identifying the true simulated drivers. For f-scLVM the factor relevance was used to rank factors. Analogous metrics were derived for all alternative methods; see Implementation details of alternative methods.

Additionally, we assessed the ability of f-scLVM to augment corrupted gene set annotations (Fig. 2c, Supp. Fig. 4). We evaluated the ability of the model to correct the false positive and false negative annotation separately.

#### Staged mouse embryonic stem cells

The set of 182 mESCs staged for the cell cycle have previously been described in ^2^. Briefly, cells were cultured in serum-free NDiff 227 medium (Stem Cells Inc.) supplemented with 2i inhibitors and sorted by cell cycle phases (G1, S G2/M) using FACS and Hoechst staining (Hoechst 33342; Invitrogen). Cells in all three cell cycle stages were profiled using the Fluidigm C1 system. We followed the pre-processing and normalization approach as previously described ^9^ and considered log-transformed and size-factor adjusted (geometric library size on endogenous genes) gene expression counts for 6,635 variable genes for analysis. Additional results show in Supp. Fig. 1c,d were obtained when considering a size-factor normalizations based on ERCC spike-ins, which retains variation in the overall amount of mRNA per cell. f-scLVM was applied using 44 gene sets derived from MSigDB (after filtering, Supp. Methods). We further applied PAGODA to the raw count data of the 182 cells using the R package scde with standard settings ^10^.

#### Zeisel et al. dataset

We analyzed log-transformed gene expression values of 3,005 single neurons sequenced using a protocol with unique molecular identifier ^16^. We followed the preprocessing and filtering steps from the primary publication, resulting in 7,097 variable genes. f-scLVM was applied using 161 annotations from the REACTOME database (after filtering, Supp. Methods), providing a high resolution annotation. Following model diagnostics steps (see *Diagnostics and f-scLVM parameter settings*), an additional 5 sparse unannotated factors were added and fit jointly with the remaining factors (Supp. Fig. 5). Residual datasets were generated by regressing out the effect of the most relevant unannotated factor (Supp. Fig. 6e-h).

#### 49,300 Retina cells

We considered the normalized, log-transformed expression values of 49,300 retina cells as described in ^21^. We considered all expressed genes, using the dropout noise model in f-scLVM to account for low sequence coverage. We considered gene sets from the REACTOME database. Because of the size of the data, we used factor pre-screening to reduce the set of factors before training, retaining 50 gene sets (Supp. Methods). To generate expression values corrected for confounding factors, we considered residual gene expression profiles, regressing out the effect of the most relevant unannotated dense factor (Supp. Table 2). Visualizations of corrected and raw expression values of six related cell types identified in the primary publication (Müller glia, astrocytes, fibroblasts, vascular epithelium, pericytes and microglia) were obtained using t-SNE.

